# mRNA-based vaccines against SARS-CoV-2 do not stimulate interferon stimulatory gene expression in individuals affected by Aicardi Goutières Syndrome

**DOI:** 10.1101/2022.05.18.492546

**Authors:** Asako Takanohashi, Mohamad-Gabriel Alameh, Sarah Woidill, Julia Hacker, Benjamin Davis, Guy Helman, Francesco Gavazzi, Laura Adang, Russell D’Aiello, Patrick Winters, Devon Cordova, Taibeen Khandaker, Houping Ni, Ying Tam, Paulo Lin, Drew Weissman, Justine Shults, Adeline Vanderver

**Author notes:** Correspondence to: Adeline Vanderver. These authors contributed equally (Asako Takanohashi, Mohamad-Gabriel Alameh).

## Abstract

The severe acute respiratory syndrome coronavirus 2 (SARS-CoV-2) poses threats to individuals with rare disease, in part because so little is known about the impact of COVID-19 infection and vaccination safety in rare disease populations. Of particular concern, given the overlap in disease manifestations and interferon dysregulation, are a group of heritable autoinflammatory conditions called type I interferonopathies. The most common of these, Aicardi Goutières Syndrome (AGS), is caused by altered nucleic acid metabolism and sensing, resulting in additional concerns surrounding the use of mRNA vaccination approaches. To determine whether mRNA vaccines induce an interferon response in AGS, we applied mRNA SARS-CoV-2 vaccines to whole blood samples and assessed internalization and interferon signaling gene expression responses to the mRNA. In all cases (11 AGS and 11 control samples), interferon signatures did not significantly increase from baseline, regardless of baricitinib treatment status in the AGS subjects, and were even decreased, when using codon optimized SARS-CoV-2 di-proline modified spike sequence (S2P). Internalization of S2P in human dendritic cells was verified by Western Blot, and in control and AGS blood cells was verified by Luciferase activity. Although numbers of tested samples in this rare disease are small, based on these findings, we suggest that COVID vaccination is unlikely to directly stimulate the interferon signaling gene expression in AGS patients via response to mRNA internalization. The *in vitro* nature of this study cannot exclude an exaggerated interferon response to spike protein production at a systemic level in individuals with a primary heritable interferonopathy. In the context of continued SARS-CoV-2 spread in the community, we do not recommend withholding vaccination in this rare disease group. However, we recommend that vaccinations for AGS patients are provided in a controlled setting with appropriate observation and used with caution in individuals with prior vaccine associated adverse events.

## Introduction

Vaccination represents a safe and effective approach to prevent and control serious, and sometimes deadly, infectious diseases [15]. The ongoing coronavirus disease 2019 (COVID-19) pandemic has required rapid development of vaccines to prevent disease, reduce mortality, and curb the ongoing crisis. Approved vaccines have been shown to be largely safe and well tolerated with demonstrated efficacy in both randomized clinical trials (RCTs) and clinical practice [32]. More recently, as mRNA vaccines have been approved for the pediatric population, individuals with rare disease remain a marginalized group in vaccination efforts due to limited safety data.

Rare genetic disorders affecting the production of interferon (IFN) are known as type I interferonopathies [9]. The type I interferonopathies include Aicardi Goutières Syndrome (AGS), a neurologic disease with early infantile-onset and clinical features caused by systemic inflammation [10, 11, 23, 24, 26]. This systemic autoinflammatory state is believed to result from poor discrimination between endogenous and exogenous nucleic acids, leading to constitutive upregulation of the IFN pathway [25]. Upregulation of IFN pathways can be assessed by measuring the levels of interferon signaling gene (ISGs) expression from whole blood [17, 27]. ISG signaling in blood samples has been used in AGS clinical trials to understand therapeutic target response [1, 34].

In viral infections, pathogen-derived nucleic acids (NAs) recognition by cell membrane, endosomal, and intracellular sensors leads to the production of type I IFN and subsequent ISG expression. IFN signaling upon SARS-CoV-2 infection is critical for disease control [7]. IFN production results from endosomal recognition by toll-like receptor (TLR)-3 and TLR-7, and components of the inflammasome including melanoma differentiation-associated protein 5 (MDA-5), retinoic acid-inducible gene I (RIG-I), Nucleotide Binding Oligomerization Domain Containing 2 (NOD2), and protein kinase R (PKR) among others coming together upon binding of single- or double-stranded RNA fragments [31]. IFN inhibits viral replication while assisting in stimulating the adaptive immune response.

mRNA-based vaccinations have been a rapid development in the clinical arsenal against SARS-CoV-2 infection [28, 35] and other infectious diseases [3]. Translation of mRNA vaccines and other mRNA based therapeutic modalities in clinical trials was enabled by both the discovery that modified nucleosides, and proper purification, improved translation and reduced the innate immune activating potential of *in vitro* transcribed mRNA, with the parallel development of efficient non-viral delivery systems [2, 20, 22, 36]. mRNA delivery has been enhanced by the addition of lipid nanoparticles (LNPs) as a capsule for the modified mRNA [22]. BNT162b (Comirnaty, Pfizer–BioNTech [35]) and mRNA-1273 (Spikevax, Moderna [5]) hereafter referred to by their commercial names, are nucleoside modified mRNA encapsulated into LNPs (mRNA-LNPs). mRNA-based vaccines are thought to provide a safer and more effective form of immunity compared to other vaccine types [14]. Studies from the development of nucleoside-modified mRNA demonstrated a lack of induction of proinflammatory cytokines and type 1 interferons in dendritic cells treated *in vitro* or after *in vivo* administration [16].

AGS is a disorder of disrupted nucleic acid metabolism that may be treated with immunosuppressive medications [13, 33, 34], making unvaccinated affected individuals at risk for severe SARS-CoV-2 infection. It is currently unclear whether mRNA-based vaccination may have the potential for detrimental innate immune activation and high levels of inflammation or whether the use of mRNA-LNP based platforms may mitigate this upregulation. To evaluate the immunostimulatory potential of these vaccines in AGS individuals, we formulated nucleoside modified mRNA into LNPs and assessed their impact on peripheral blood monocyte cells in whole blood collected from individuals affected with AGS versus controls. We measure ISGs, a measure of IFN activation used in clinical trials, to assess the IFN response *in vitro* after addition of mRNA-LNPs.

## Materials and Methods

### Cohort identification and sample collection

Disease and control samples were collected under a leukodystrophy associated biorepository protocol, the Myelin Disorders Biorepository and Natural History Study, approved at Children’s Hospital of Philadelphia (IRB 14-011236). Individuals were identified as having either AGS (Disease Cohort) based on molecular identification of one of the AGS-related genes (*TREX1* [n=1], *RNASEH2B* [n=3], *SAMHD1* [n=3], *ADAR1* [n=3], or *IFIH1* [n=1]; no patients variants in *RNASEH2A, RNASEH2C, RNU7-1* and *LSM11* were identified) or as having a non-AGS related leukodystrophy or their unaffected parents (Control Cohort) (Table 1). Absence or presence of treatment with the Janus Kinase Inhibitor, baricitinib, was noted. In the Disease and Control Cohorts, samples were drawn during a venipuncture immediately into EDTA containing tubes and aliquoted into 1 mL aliquots. Whole, unprocessed blood was incubated with mRNA-LNP vaccines, and associated controls as detailed below.

**Table 1.** Cohort demographics and ISG scores.

| Study ID | Diagnosis | Affected gene | Age (y) | Baricitinib Treatment | Baseline ISG scores | Nucleoside Modified mRNA-LNP (30$\mu\text{g}/\text{mL}$) ISG scores |
| --- | --- | --- | --- | --- | --- | --- |
| **Controls** |  |  |  |  |  |  |
| LD_1252.0 | AxD | *GFAP* | 17.6 | No | -0.27 | -0.61 |
| LD_1255.0 | AxD | *GFAP* | 11.2 | No | -0.07 | -0.43 |
| LD_1980.0 | TUBB4A | *TUBB4A* | 13.3 | No | -0.29 | -0.49 |
| LD_0313.0 | TUBB4A | *TUBB4A* | 20.3 | No | 0.66 | -0.43 |
| LD_2261.0 | TUBB4A | *TUBB4A* | 3.5 | No | 0.03 | -0.45 |
| LD_1504.0 | Unknown | multiple | 6.1 | Yes | -0.75 | -0.73 |
| LD_2261.1 | Parent | N/A | 32.1 | No | -0.47 | -0.74 |
| LD_2261.2 | Parent | N/A | 29.4 | No | 0.17 | -0.13 |
| LD_0313.1 | Parent | N/A | 44.4 | No | 0.89 | 0.45 |
| LD_0313.2 | Parent | N/A | 58.0 | No | -0.01 | -0.47 |
| LD_1706.1 | Parent | N/A | 33.6 | No | -0.39 | -0.59 |
| **AGS Patients** |  |  |  |  |  |  |
| LD_1185.0 | AGS | *TREX1* | 5.4 | Yes | 16.61 | 10.78 |
| LD_2100.0 | AGS | *RNASEH2B* | 1.2 | No | 2.21 | 1.71 |
| LD_1574.0 | AGS | *RNASEH2B* | 10.8 | Yes | 4.02 | 1.19 |
| LD_1177.0 | AGS | *RNASEH2B* | 6.8 | Yes | -0.99 | -0.94 |
| LD_1631.0 | AGS | *SAMHD1* | 17.1 | No | 21.76 | 11.95 |
| LD_1471.0 | AGS | *SAMHD1* | 10.6 | Yes | -0.47 | -0.60 |
| LD_1254.0A | AGS | *SAMHD1* | 25.0 | Yes | 41.48 | 23.16 |
| LD_1669.0 | AGS | *ADAR1* | 7.5 | Yes | 10.43 | 3.54 |
| LD_2096.0 | AGS | *ADAR1* | 2.2 | No | 37.34 | 11.51 |
| LD_1034.0 | AGS | *ADAR1* | 8.0 | Yes | 16.02 | 10.10 |
| LD_0940.0 | AGS | *IFIH1* | 8.2 | Yes | 18.6 | 10.23 |

AxD - Alexander Disease; TUBB4A - *TUBB4A*-related leukodystrophy; AGS - Aicardi Goutières Syndrome

### Cohort Identification for patient reported outcomes

In a distinct cohort from the above cohort collected for samples, the Aicardi Goutières Syndrome Americas Association (AGSAA) queried patients enrolled in an IRB approved patient registry, Luna (Luna software, version 2022 of LunaPBC, Inc. Copyright 2022, Genetic Alliance), with a standard series of questions around vaccine and COVID-19 vaccine experience. Patients or their care givers entered responses to the questionnaire. Individuals were identified as having molecular identification of one of the AGS-related genes (*TREX1* [n=2], *RNASEH2B* [n=7], *RNASEH2C* [n=1], *SAMHD1* [n=7], *ADAR1* [n=2], *IFIH1* [n=3], *RNU7-1* [n=2]); no patients variants in *RNASEH2A* and *LSM11* were identified). Genotype data was unknown or unavailable for 11 respondents.

### Production of the mRNA encoding the di-proline stabilized SARS-CoV-2 spike protein and the firefly luciferase protein

Codon optimized SARS-CoV-2 di-proline modified spike sequence (S2P), and firefly Luciferase (fLuc) sequence were cloned into an mRNA production plasmid (optimized 3’ and 5’ UTR and containing a 101 polyA tail), *in vitro* transcribed in the presence of UTP (unmodified) or in the presence of N1-methylpseudouridine modified nucleoside (N1-m*ψ*, modified), co-transcriptionally capped using the CleanCap™ technology (TriLink) and cellulose purified to remove dsRNA [6]. Purified mRNA was ethanol precipitated, washed, re-suspended in nuclease-free water, and subjected to quality control (electrophoresis, dot blot, endotoxin content and transfection into human DCs). mRNA sequence used in this study is identical to the one used in the Pfizer/BioNTech and Moderna vaccines except for the 3’ and 5’ UTRs. mRNA was stored at -20°C until use.

### Production of empty LNPs and mRNA-LNP vaccines

mRNA loaded LNPs, or vaccines, were formulated using a total lipid concentration of 40mM as previously described [3]. The ethanolic lipid mixture comprising ionizable cationic lipid, phosphatidylcholine, cholesterol and polyethylene glycol-lipid was rapidly mixed with an aqueous solution containing cellulose-purified UTP or N1-m*ψ in vitro* transcribed mRNAs to generate the unmodified mRNA-LNP vaccine and the nucleoside modified mRNA-LNP resembling the clinical vaccine Comirnaty, respectively [21]. The LNP formulation used in this study is proprietary to Acuitas Therapeutics; the proprietary lipid and LNP composition are described in US patent US10,221,127. Empty particles were prepared as described above without the addition of mRNA in the aqueous phase. RNA-loaded and empty particles were characterized and subsequently stored at -80°C at an RNA concentration of 1 *μ*g/*μ*L (in the case of loaded particles) and total lipid concentration of 30 *μ*g/*μ*L-1 (both loaded and empty particles). The mean hydrodynamic diameter of mRNA-LNPs was 80 nm with a polydispersity index of 0.02-0.06 and an encapsulation efficiency of 95%. Two or three batches from each mRNA-LNP formulations were used in these studies.

### *In vitro* mRNA expression to demonstrate the bioactivity of mRNA-LNP vaccines

Expi293F cells were transiently transfected with 3*μ*g mRNA encoding the nucleoside modified S2P using Lipofectamine messenger Max™ or mRNA-LNP. Transfected cells were harvested 48 hours post-transfection, washed once, resuspended in FACS buffer (1X PBS, 2% FBS, 0.05% sodium azide) at 1×10^6^ cell/ml, and stained with 10*μ*g/mL D001 antibody (SinoBiological) in U-bottom 96 well plates for 30 minutes at 4°C. Cells were washed once and incubated with a PE-labelled goat anti-human IgG secondary antibody (Thermo Fisher Scientific 12-4998-92) at a final concentration of 2.5*μ*g/mL of 30 minutes at 4°C. Dead cells were stained with amine reactive Live/Dead fixable aqua dead cell stain kit for 5 minutes, washed, and resuspended in 200*μ*L 1% paraformaldehyde containing FACS buffer. Flow cytometric data was acquired using an LSRII and data analyzed using FlowJo 10.2.

### Generation of human DC from blood and detection of IFN-*α* following transfection with mRNA-LNPs

Blood derived monocytes were isolated at the University of Pennsylvania Human Immunology Core (HIC) and differentiated into monocyte derived dendritic cells (MoDCs) using 50ng/mL human IL-4 and 100ng/mL GM-CFS in R10 media. Monocytes were differentiated for 3 days, and MoDCs counted, and seeded at 1 million cells/well. mRNA-LNP were added to each well at 0.3*μ*g/well, and the supernatant collected 24 hours post transfection to measure IFN-*α* using the Human IFN-*α* Pan specific ELISA kit (MabTech). MoDCs were lysed using RIPA buffer supplemented with protease inhibitors, clarified using centrifugation at 10,000 g for 1 minute, and total protein content was measured using the microBCA kit (Pierce). 10*μ*g total protein was denatured at 65°C in the presence of 1 X Laemmli buffer and loaded on a 14% SDS–PAGE gel. Electrophoresis was performed for 1 hour at 100 V, and the gel transferred to a Hybond PVDF membrane using the i-blot system (Thermo Fisher scientific). Membranes were blocked in 5% non-fat milk for 1 hour, washed twice with TBS-T, incubated with rabbit anti-SARS-2-S at 1:2000 (Sino Biological 40591-T62) in 1% non-fat dry milk for 1 hour, washed twice in TBS-T and incubated in the presence of a goat HRP conjugated anti-rabbit secondary IgG antibody (1:10000; Abcam 6721). Membranes were revealed using the Amersham ECL prime reagent (GE healthcare) and imaged on an Amersham A600 system. Membranes were stripped and re-blotted in the presence of an anti-GAPH antibody as a loading control.

### Internalization of mRNA-LNPs encoding the firefly luciferase protein

Whole blood samples containing 10% RPMI-1640 medium were incubated at 37°C for 6 hours with nucleoside modified mRNA-LNP, which is mutated to express whole spike protein with no-cleavage. Blood samples were collected and PBMCs were purified using HISTOPAQUE-1077 (Sigma) followed by PBS wash. PBMCs were lysed in 50*μ*L of RIPA buffer and assessed using the Promega One Glo Luciferase assay.

### Interferon Signaling Gene (ISG) Expression from whole blood samples

One mL of whole blood collected into EDTA blood tube (BD) was mixed with 10% RPMI-1640 and plated 1.1 mL per well in a 12-well plate. mRNA-LNP vaccines, empty LNPs (eLNP), and naked mRNA was added to each blood containing well, and the plate was incubated 6hr in a 5% CO2 incubator at 37°C. At the end of the incubation, samples were mixed with a 3 mL of PAXgene RNA stabilization reagents (PreAnalytix) to stabilize RNA and further incubated at room temperature for up to 24hr to allow complete RBC lysis. RNA stabilized blood samples were kept at -80°C until use. RNA was purified using PAXgene blood RNA kit (Qiagen) following manufacturer recommendations. Total RNA concentration was determined using the Quant-iT RNA HS Reagent (Thermo Fisher Scientific) and 200ng of RNA per sample was used to measure gene expression as detailed below.

NanoString nCounter® analysis approaches have been previously described for the measurement of ISG expression [17]. After hybridization at 65°C for 16h, nCounter “Elements” sample runs were immediately processed using the “High Sensitivity” protocol option on the nCounter FLEX Prep Station and tagged RNA numbers of each gene were counted on nCounter FLEX Digital Analyzer using maximal data resolution (555 fields of view). Data were processed with nSolver software (NanoString Technologies, Seattle, WA), which included assessment of quality of the runs using Nanostring methodologies of assay validity and within run standards for ISG oligo sequences.

For ISG scores, RNA counts are normalized to 4 housekeeping genes (*ALAS1, HPRT1, TBP*, and *TUBB*), then used to calculate z-scores in Stata (Collage Station, Texas) as previously published.11 The median of the z-scores of six ISGs based on the gene list provided by Rice et al. (*IFI27* + *IFI44* + *IFI44L* + *ISG15* + *RSAD2* + *SIGLEC1*) [10, 25, 27] was used to generate an ISG score, as previously used in AGS related studies [4, 34]. To generate the heatmap, for each ISG signaling gene comparisons, two normalization processes were performed on the raw RNA counts to generate normalized counts: (1) Six internal positive control sequences that are not native to any known organism, are run with each sample. The geometric mean is calculated for these positive control sequences at each sample run, and a scaling (normalization) factor is calculated for each sample to equilibrate each count for the respective gene to the geometric means of the positive controls. (2) A second normalization (content normalization) is performed by using the same process by applying a scaling factor that “normalizes” the geometric mean of the housekeeping genes (*ALAS1, HPRT1, TBP*, and *TUBB*) . These normalized counts are used to compare each gene. A heatmap was generated using GraphPad Prism (GraphPad Software, San Diego, CA).

#### Statistical Analysis

We treated the ISG scores for each vaccine type as a cluster of measurements on each subject. Our goal was to evaluate the effect of treatment on ISG scores and to compare the effect between AGS and non-AGS subjects. We first evaluated the impact of treatment on change from no treatment in all subjects (Table 2) by fitting quasi-least squares (QLS) regression models [8] with the following variables: an indicator variable for each of the 4 vaccine types (with untreated as the reference category); an indicator variable for control (versus AGS); and baseline ISG (untreated). To account for the correlation among the multiple ISG scores per subject, we fit the exchangeable correlation structure with the robust sandwich covariance estimator of covariance of the regression parameter. After fitting the model, we used the lincom procedure in Stata 17 to estimate the difference in ISG scores (with 95% confidence intervals [CI] and p-value for test that the difference is zero). We did not fit the unstructured correlation matrix for this analysis because application of the QLS matrix for QLS requires an equal number of measurements per subject; 13 subjects had 5 vaccine types, while 9 subjects had 7 vaccine types. Generalized estimating equations (GEE) allows for an unequal number of measurements per subject for the unstructured matrix, but GEE failed to converge for this structure.

**Table 2.** Estimated Difference in ISG Scores between Untreated and Treated Samples for each vaccination type.

| Vaccine Type | Estimated Difference | 95% CI | p-value |
| --- | --- | --- | --- |
| Low-dose Unmodified mRNA vaccine (n=22) | 5.39 | (3.09 to 7.70) | <0.0005 |
| High-dose Unmodified mRNA vaccine (n=22) | 15.37 | (10.83 to 19.91) | <0.0005 |
| Empty LNP (eLNP [n=22]) | -0.58 | (-1.35 to 0.19) | 0.142 |
| Nucleoside Modified mRNA Vaccine (n=22) | -3.21 | (-5.20 to -1.22) | 0.002 |
| Naked Unmodified Vaccine (n=9) | 0.04 | (-0.71 to 0.79) | 0.918 |
| Naked Modified Vaccine (n=9) | -0.15 | (-0.88 to 0.59) | 0.697 |

We next evaluated the impact of treatment on change from no treatment (Table 3) while also allowing the change from no treatment to vary between AGS and non-AGS samples. We accomplished this by fitting QLS regression models on change from no treatment with the following variables: an indicator variable for each of the 4 vaccine types (with untreated as the reference category); an indicator variable for control (versus AGS); baseline ISG (no treatment); and a control by treatment type interaction term. Naked unmodified and naked modified could not be included in this analysis because there was only one AGS sample for these vaccine types. Due to the limited sample size, we included the interaction terms one at a time, in separate models for each treatment type. To account for the correlation among the multiple ISG scores per subject, we fit the exchangeable and unstructured correlation matrices. We evaluated the fit of the exchangeable and robust structures using several criteria and selected the structure with the best fit. We then fit the model for the best fitting correlation structure. After fitting the model, we used the lincom procedure in Stata 17 to estimate the difference in ISG scores (with 95% confidence intervals [CI] and p-value for test that the difference is zero) between treatment types.

**Table 3.** Estimated Difference in ISG Scores in AGS and non-AGS samples between Untreated and Treated Samples for each vaccination type.

| Vaccine Type | Estimated Difference (ISG for vaccine minus baseline) with 95% CI for AGS samples (n=11) | Estimated Difference (ISG for vaccine minus baseline) with 95% CI for non-AGS samples (n=11) | p-value for test for interaction between vaccine type and AGS |
| --- | --- | --- | --- |
| Low-dose Unmodified mRNA vaccine (n=22) | 7.24 (95% CI = 3.48 to 11.01) | 3.54 (95% CI = 1.80 to 5.29) | 0.0609 |
| High-dose Unmodified mRNA vaccine (n=22) | 18.48 (95% CI = 12.40 to 24.57) | 12.26 (95% CI = 6.11 to 18.41) | 0.1504 |
| Empty LNP (eLNP [n=22]) | -1.75 (95% CI = -3.81 to 0.31) | 0.60 (95% CI = -0.83 to 2.03) | 0.1579 |
| Nucleoside Modified mRNA Vaccine (n=22) | -6.87 (95% CI = -10.63 to -3.11) | 0.46 (95% CI = -1.06 to 1.97) | 0.0024 |

## Results

### Validation of internalization, immunogenicity and expression of SARS-CoV-2 di-proline modified spike sequence (S2P) in human dendritic cells in culture and Expi293F cells

mRNA-LNP encoding the di-proline modified spike protein were transfected into human blood derived dendritic cells and commercially available high density human embryonic kidney cell line (Expi293F cells) at a dose of 0.3*μ*g/million cells. Western blot analysis demonstrated that the spike protein was efficiently translated 24 hours post transfection into human blood derived DCs with an expected band around 165 kDa (Figure 1A). The specificity of the spike protein was demonstrated by the lack of expression in the empty LNP control. Analysis of the supernatant demonstrated that nucleoside modified mRNA-LNP is not immunogenic with no detectable levels of TNF-*α*. In contrast, and as expected, uridine containing mRNA-LNP vaccines induced high levels of TNF-*α* (Figure 1B). Flow cytometry analysis of cells transfected with the mRNA-LNP vaccine showed that expression of the di-proline modified spike protein could be detected at high levels on the cell surface (Figure 1C). Detection of the protein using western blot or flow cytometry in whole blood was difficult to achieve. In order to demonstrate that these LNPs are able to enter cells and express their cargo following their addition into blood, we used a luciferase reporter mRNA for improved sensitivity. Blood from AGS patients and control subjects was transfected with either Luc mRNA-LNP or empty for 6 hours, and PBMCs were purified, lysed and subjected to the luciferase assay. Luciferase expression was detected at least 1000 fold higher than control (or background) (Figure 1D) indicating that mRNA-LNPs are able to transfect target cells in blood.

**Figure 1.**
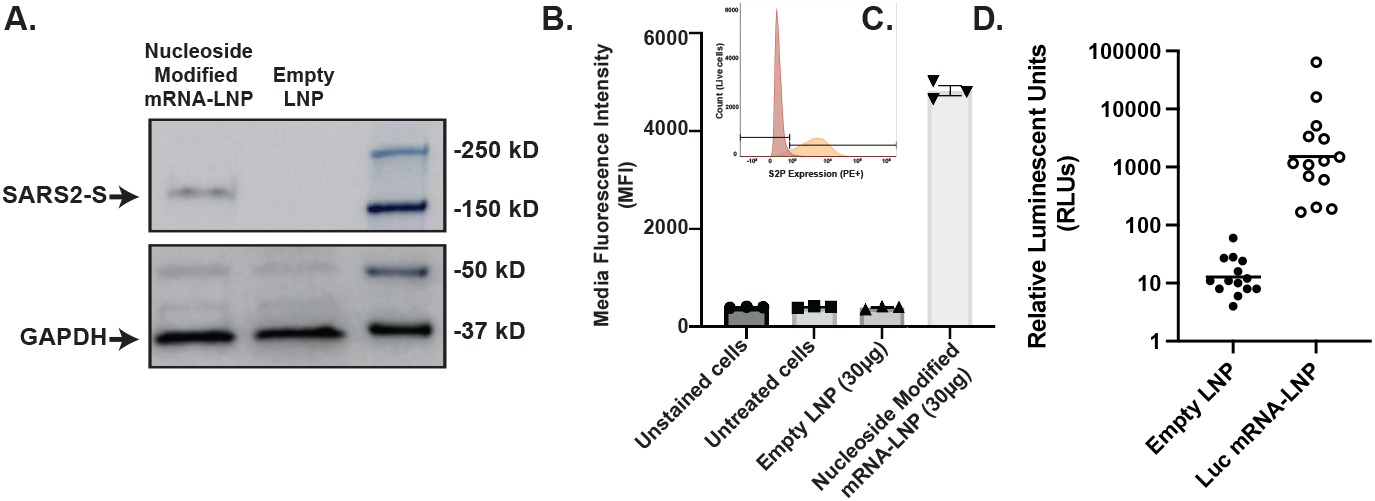
Validation of the production of di-proline stabilized spike (S2P) following mRNA encapsulation in LNPs. mRNA was produced, encapsulated into LNPs to form the vaccines, and transfected in multiple cell types to validate expression. (A) Expression of the S2P protein by western blot following transfection of human dendritic cells. (B) Expi293F cells were cultured in the presence of 30ug of S2P mRNA LNP or eLNP for 48 hours. (C) Cells were harvested and stained using the anti-Sars-CoV-2 spike D10 monoclonal antibody, and the expression of the S2P on the surface of the cells measured using flow cytometry. (D) Expression of Luc in AGS and control patient blood 24 hours post transfection with 5μg Luc mRNA-LNP.

### Interferon Stimulatory Gene (ISG) scores in AGS and control subjects with and without SARS-CoV-2 mRNA vaccine treatment

ISG scores are known to be increased when the IFN signaling pathway is stimulated, as in AGS. Most AGS patients (8/11) demonstrate an elevated ISG score (*>*1.96) at baseline compared to controls (n=11, average ISG scores: -0.05±0.48) regardless of treatment with baricitinib or not (Table 1). After treatment with the unmodified mRNA-LNP vaccine (5 and 30*μ*g/mL), an increase was seen in ISG scores in both control and AGS patient cells with a significant increase relative to baseline in non-AGS controls at 30*μ*g/mL (p=0.0131) (Figure 2). However, after the nucleoside modified mRNA-LNP vaccine is applied (30*μ*g/mL) there is no increase in ISG scores (Table 2, Figure 2). Empty LNP (eLNP) alone either did not change or decrease ISG expression. It should be noted that unmodified mRNA-LNP increases specific IFN signature genes (*CXCL10, IFIT1, IFIT2, ISG15, MX, RSAD2*) in a dose dependent manner (Figure 3) but no such effect is seen using modified mRNA-LNP. The nucleoside modified mRNA-LNP vaccine does not increase these signature genes. We found statistically significant changes in expression of *IFI27* (p=0.021), *ISG15* (p=0.005), *OASL* (p=0.005), and *SOCS1* (p=0.05) when comparing treatment with the between the nucleoside modified mRNA-LNP and eLNP (Figure 3). We did not find any statistically significant changes in expression of other RNA sensing genes (*MDA5/IFIH1, OAS1, OAS2, OAS3, PKR/EIF2AK2, RIG-I/DDX58*) between baseline and treatment across any condition (Figure 3). Of note, AGS versus non-AGS individuals were not significantly different across all vaccine types, except for the nucleoside modified mRNA-LNP vaccine, in which ISG expression was significantly decreased.

**Figure 2.**
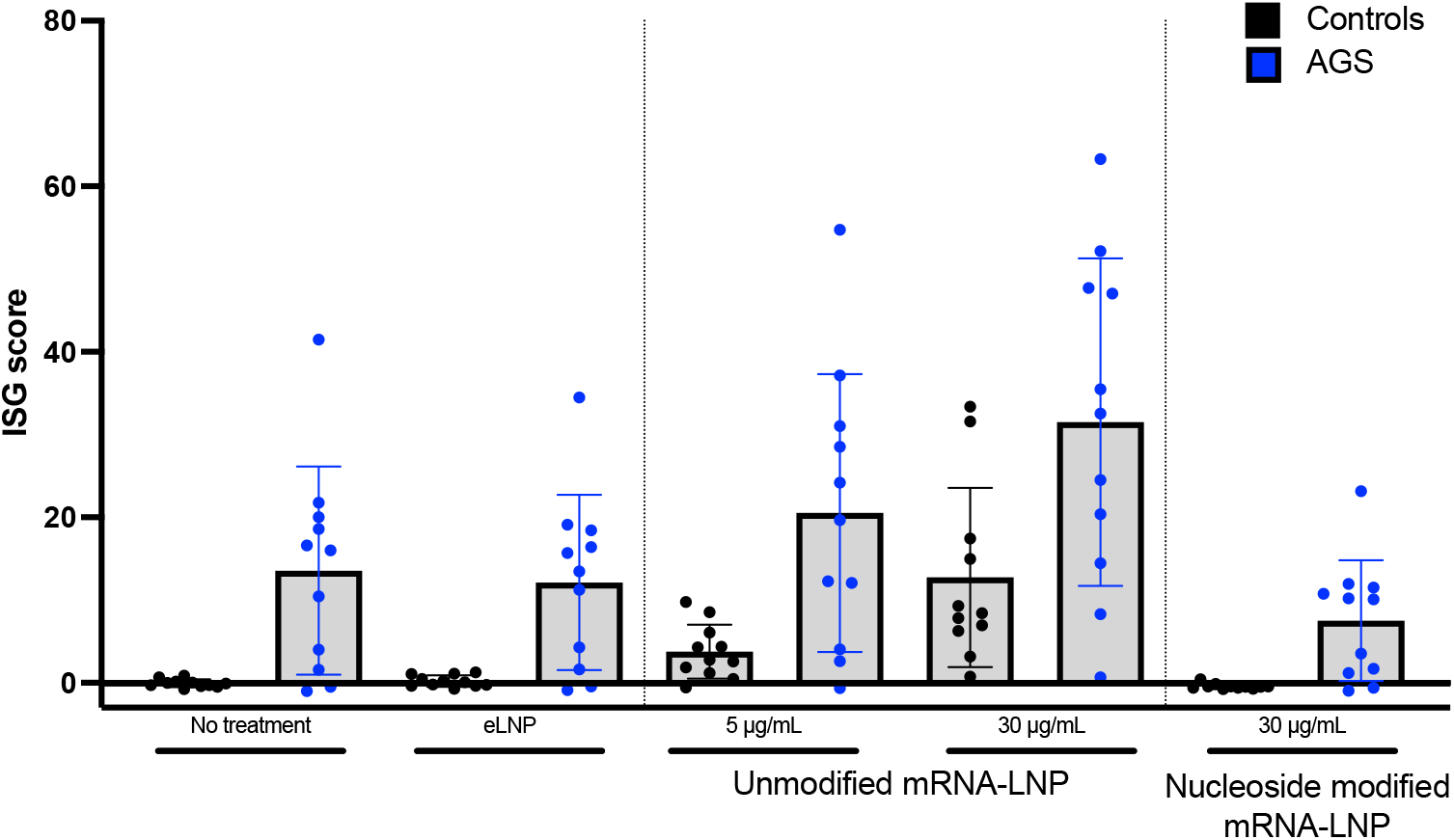
Measurement of interferon signaling gene expression scores after incubation with vaccine formulations. Interferon signaling gene (ISG) scores were assessed for all patients and controls following *in vitro* vaccine treatment. Most AGS patients (8/11) demonstrate an elevated ISG score (*>*1.96) at baseline compared to controls (n=11) regardless of treatment with baricitinib or not. The empty LNP (eLNP) did not affect ISG scores while they were elevated upon treatment with the and at both doses of the unmodified mRNA-LNP vaccine (5 and 30μg/mL) in both control and AGS patient cells. There was no increase in ISG scores following treatment with the nucleoside modified mRNA-LNP.

**Figure 3.**
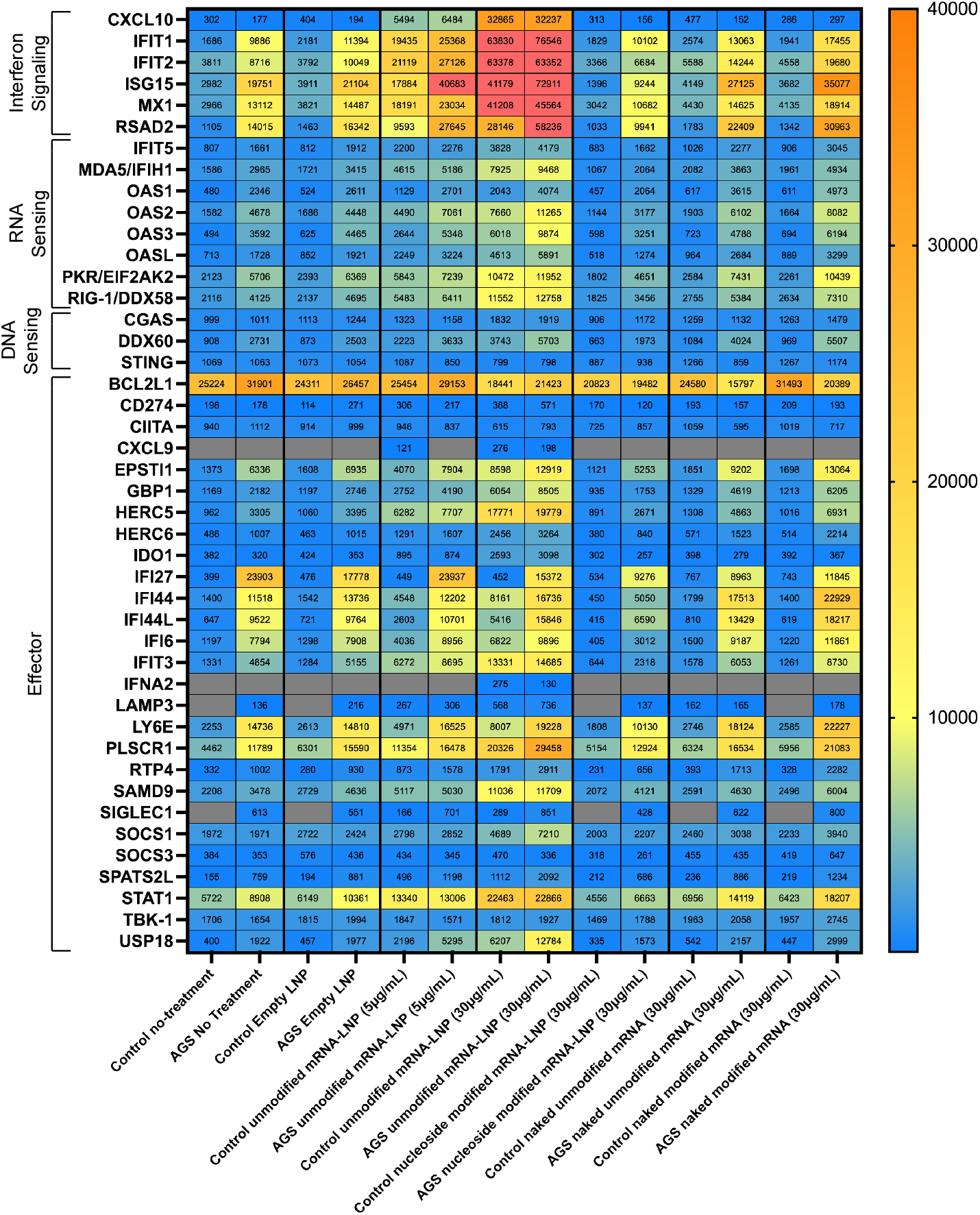
Breakdown of interferon signaling gene expression scores after incubation with vaccine formulations compared to empty LNP administration. Assessment of gene expression changes in AGS patients versus controls across conditions and categorized based on role in DNA and RNA sensing. Interferon signaling gene (ISG) expressions were increased at baseline for all patients and following *in vitro* vaccine treatment across all conditions, albeit more significantly increased upon treatment with unmodified mRNA vaccination, similar to controls. RNA sensing genes are induced only upon treatment with unmodified mRNA LNP vaccines, and similarly effector molecules are induced following treatment with UTP containing mRNA. There is no elevation of DNA sensing genes across conditions.

### Patient reported experience with COVID-19 infection and vaccination

A total of 35 individuals reported experience with COVID-19 infection and vaccination (Figure 4). 12 individuals experienced a patient reported COVID-19 infection; of these individuals 3 indicated that subsequent to infection they had long term impact to their health. Of the 12 individuals who experienced a known COVID-19 infection, 8 were not vaccinated, 2 were vaccinated and 2 had an unknown vaccination status. 16 individuals have not experienced a known COVID-19 infection. Of these 16 individuals, 8 individuals received a COVID-19 vaccination course. Among all 12 vaccinated individuals, mRNA vaccines were used in 10 individuals; 2 individuals who were 18 or older received Johnson & Johnson vaccination. Of the vaccinated individuals, 9 individuals had no side effects to vaccination, and 3 individuals had expected side effects (fever, muscle pains, headache and fatigue). No individual had unexpected side effects to vaccination.

**Figure 4.**
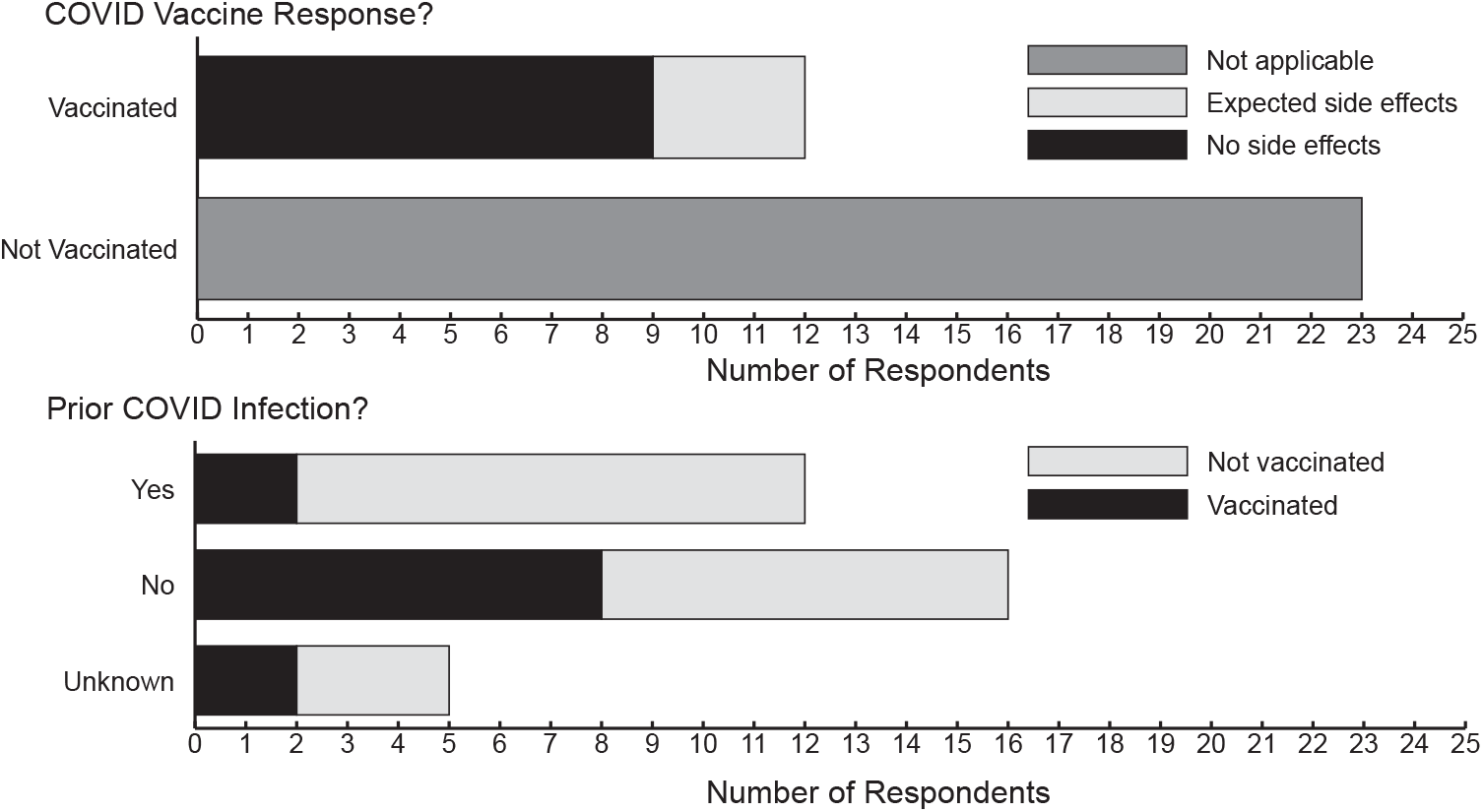
COVID Infection by Vaccination Status in Patient Reported Survey. Surveys were conducted by the AGSAA of their membership. 35 members reported vaccination status and 30 members reported infection status.

## Discussion

AGS and related heritable interferonopathies have unregulated IFN production, which can worsen in the context of infection or other immune triggers [1]. In addition, several AGS related genes are involved in the detection (sensing) and the regulation of endogenous nucleic acids [11, 12, 23, 24, 26]. This has led to concern that nucleic acid based vaccines such as the FDA approved Comirnaty [35] and Spikevax [5] mRNA-based vaccines could result in further dysregulation of IFN signaling and has led to hesitancy around vaccine adoption in this rare disease population.

Our experiments do not suggest an increase in IFN signaling, as measured by the calculation of an ISG score, after internalization of the clinical S2P in a LNP (Table 1, Figure 1). Although unmodified mRNA vaccines increase ISGs in both control and AGS samples, the internalization of the nucleoside modified mRNA vaccine representing the clinical vaccine significantly decreases ISGs relative to untreated samples in both control and AGS samples. It should be noted that the SARS-CoV-2 spike protein itself has been shown *in vitro* to result in decreased production of type 1 IFN [30], but that SARS-CoV-2 overall has a complex interaction with IFNs, likely associated with tissue specificity and disease severity [18, 29, 31]. It is promising that the expression of RNA sensing genes is upregulated only in the presence of unmodified mRNA vaccine, as expected. However, it remains unclear if the decreased production *in vitro* of ISGs by cultured cells after treatment with the nucleoside modified mRNA vaccine may not represent the complexity of vaccine response at the organism level in AGS affected individuals.

The elevated ISGs seen on *in vitro* treatment of AGS patient cells with unmodified mRNA vaccines is of particular concern. The HERALD phase 2b/3 clinical trial using a chemically unmodified mRNA vaccine, named CVnCoV, demonstrated only 53% efficacy against SARS-CoV-2 compared to the 95% and 94% efficacy of the Comirnaty [35] and Spikevax [5] vaccines, respectively [19]. Importantly, in this trial the frequency of adverse events was more frequently reported in CVnCoV recipient population than in a placebo group in healthy participants despite a reduced dose (12μg) being used to immunize participants [19]. We observed elevated ISGs and elevation in RNA sensing genes upon administration of an unmodified mRNA LNP vaccine in both AGS and control patient cells, demonstrating the immunogenicity of this formulation. We suggest that the use of mRNA-based vaccinations overall in AGS patients should be limited to nucleoside modified based formulations and the development of other unmodified mRNA LNP vaccine formulations should take this into account.

It is important to note that the safety and effectiveness of COVID-19 vaccines in those with AGS has not been studied. It is however unlikely that such a clinical trial will occur in a rare disease population. Because of limited information, our guidance is based on this limited *in vitro* data. Although a systemic autoinflammatory response to the spike protein is likely and may increase ISG signaling, it is likely that this will not be as great as the autoinflammatory response seen in the context of an actual infection. Thus, in the context of continued COVID-19 infections in the community, however, we are cautiously recommending that AGS affected individuals consider being vaccinated with one of the United States Food and Drug Administration approved mRNA vaccines, as applicable for their ages in the general population, unless individuals have documented vaccine related adverse events.

## Acknowledgments

We thank the patients and their families, and the AGSAA for their contributions to this data. JS, AV, AT, FG and LA received funding from U54TR002823 (NCATS and NINDS) and U01 NS106845 (NINDS), as well as the Children’s Hospital of Philadelphia Research Institute. LA received support from NINDS K23NS114113. The patient reported data for this study was collected using Luna software, version 2022 of LunaPBC, Inc.Copyright © 2022 Luna. Restrictions apply to the availability of the data which were used under license for this study. Data are available from the authors with the permission of the study participants in Luna.

## Data availability

The data that support the findings of this study are available on request from the corresponding author. The data are not publicly available due to privacy or ethical restrictions.

